# A High-throughput Sequencing Screen to Identify Apomixis in Plants

**DOI:** 10.1101/2025.11.12.688100

**Authors:** Charity Z. Goeckeritz, Václav Polcar, Benjamin Gutierrez, Tomáš Urfus, Alex Harkess

## Abstract

In the past decade, plant biologists have made several major discoveries pertaining to the genetic basis of apomixis (clonal propagation by seed) that have shown promise in preserving high-value hybrid rice and sorghum genotypes. This progress was made possible by foundational gene discovery efforts in model species and natural apomicts, but pleiotropic obstacles still limit its broad agricultural adoption, especially in eudicots. Thus, it follows that investigations of novel apomicts will lead to the development of new molecular tools for plant breeding. The three most common ways to identify clonal seed production are cytoembryological analyses, genome sequencing to compare the DNA sequence of mother and child, and flow-cytometry seed screens. While flow-cytometry has been the dominant method for more than two decades, it requires specialized equipment, expertise, and occasional troubleshooting. Here we developed a method using short-read skim-seq at moderately low sequencing depths (3X and 6X) to screen diverse *Malus* genotypes maintained in a USDA germplasm collection for clonal seed production. By sequencing 55 genotypes and 1,216 of their embryos, our screening-by-sequencing method found 17 previously undescribed apomictic genotypes. Five additional apomictic genotypes were detected with flow cytometry data and using internal sequencing controls. This skim-seq embryo screening-by-sequencing method is a relatively low-cost, rapid method for detecting apomictic genotypes in diverse plant germplasm, and when used thoughtfully in conjunction with flow cytometry, provides a new way to visualize the genetic outcomes of sexual and asexual reproduction in plants.

## INTRODUCTION

Apomixis, or clonal propagation via seed, is a complex reproductive process that has evolved many times in Angiosperms (Hojsgaard et al. 2014). This convergent trait has been documented in several hundred genera, including those of direct economic value (e.g., *Rubus*, *Malus*, *Cucumis*) and crop wild relatives (e.g., *Tripsacum* for maize) (Crane and Thomas 1939; Sax 1959; Zagorcheva Maritsa 1987; Grimanelli et al. 1998). Identifying the genes underlying apomixis has been a highly sought-after goal for plant biologists for decades, as these genes have the potential to revolutionize plant breeding. Not only do the genes controlling natural and synthetic apomixis allow for the preservation of valuable genotypes, but they are also being investigated for their capacity to create haploids and to strategically engineer heterotic polyploids (Qi et al. 2023; Skinner et al. 2023; Ren et al. 2024; Wang et al. 2024).

The field has had several major breakthroughs in the past decade with the characterization of *BABYBOOM* (*BBM*) in *Cenchrus squamulatus* (previously *Pennisetum squamulatum*) and *PARTHENOGENESIS* (*PAR*) in *Taraxacum officinale*, both of which are sufficient to induce parthenogenesis (the spontaneous development of an embryo in the absence of fertilization) in gametophytic apomicts (Conner et al. 2015; Underwood et al. 2022). In combination with *MiMe* (*Mitosis instead of Meiosis*) and CRISPR technology, these genes have made synthetic apomixis highly efficient in rice and its agricultural application is now on the horizon for this crop (Khanday et al. 2019; Wang et al. 2019; Vernet et al. 2022; Wei et al. 2023; Song et al. 2024; Huang et al. 2025). More recently, researchers also successfully engineered highly penetrant synthetic apomixis in two sorghum hybrids (Simon et al. 2025). Still, the applicability of *MiMe* to other systems remains unclear, especially in eudicot species. Several works have already demonstrated alternative genes to *SPO11*, *REC8,* and *OSD1* are necessary to abolish meiosis in other plants while minimizing pleiotropic effects, likely due to gene expansion, contraction, and the functional divergence of these homologs throughout evolution (Qian et al. 2024; Wang et al. 2024; Pang et al. 2025). However, the prevalence of apomixis in flowering plants means ample opportunity to discover new tools for plant breeding (Hojsgaard et al. 2014; Goeckeritz et al. 2024).

The discovery of apomixis genes has been slowed by a number of factors. Traditionally, gene discovery requires the generation of segregating populations, yet apomixis is partly defined by the absence of meiosis and recombination. While the vast majority of natural apomicts are facultative, meaning sexual reproduction still occurs at a rate that is dependent on genotype and environment (Liu et al. 2014; Klatt et al. 2018; Soliman et al. 2021), fine mapping these genes is still hindered by long generational times, polyploidy, distorted segregation, and potentially large hemizygous, non-recombining regions (Ozias-Akins et al. 1998; Van Dijk et al. 2020). Fortunately, advancing genomic technologies are beginning to offer alternative approaches to complement conventional ones. Wang *et al*. (2022) used resequencing data of diverse genotypes to fine map *RWP-RK* genes in *Citrus* and *Fortunella*, and Yadav *et al*. (2023) identified an orthologous gene in *Mangifera*. This gene had previously been confirmed in *Citrus* to induce somatic embryogenesis (Shimada et al. 2018; Wang et al. 2022). Notably, both examples mapped genes in diploids exhibiting sporophytic apomixis, which tends to segregate normally as a single dominant trait. Genomic methods will be especially impactful in perennial species with loci embedded in larger, non-recombining regions of the genome (e.g., aposporic gametophytic apomicts) by taking advantage of ancestral recombination to narrow in on gene candidates. These studies also underscore the enormous value of maintaining diverse, living collections and developing international collaborations to advance our understanding of the evolution and functional genomics of plant reproductive biology.

Given the availability of such germplasm in the adult phase, there is still the matter of screening genotypes for apomixis en masse. Several methods have been developed to screen for apomixis, including microscopy techniques to observe megagametogenesis and embryology, pollen exclusion and pistil decapitation for autonomous apomixis, the flow cytometric seed screen (FCSS), and the comparison of a variety of genetic markers such as isozymes, microsatellites, random amplification of polymorphic DNAs, and more (summarized in (Hojsgaard and Pullaiah 2022)). By far, FCSS is the preferred method to distinguish apomixis, sexuality, and intermediate forms of reproduction as it is moderate-throughput and ploidy agnostic (Matzk et al. 2000; Krahulcová and Rotreklová 2010). However, the method can be uninformative or technically challenging in specific species, such as pseudogamous apomicts with uninucleate maternal endosperm contributions or species with diminished endosperm (Matzk et al. 2000; Dobeš et al. 2013). FCSS also requires specialized equipment and expertise, which can become prohibitively expensive when outsourced as a service. Thus, it would be prudent to develop other high-throughput and low-cost methods for screening apomixis to broaden accessibility and increase the pace of gene discovery.

Sequencing-based approaches are becoming an increasingly viable option as DNA extraction has the potential to be automated, library preparation kits now allow for the multiplexing of hundreds of samples, and sequencing costs continue to decline. Here we develop a novel low-cost and high-throughput embryo sequencing screen to detect apomixis in apple (*Malus*, Rosaceae). We selected apple for the development of this screen given its agronomic importance and the availability of diverse germplasm maintained by the USDA in the adult phase. Wild *Malus* species native to eastern Asia and North America have previously been documented to exhibit aposporic apomixis (Sax 1959; Kron and Husband 2009; Dickinson 2018; Hojsgaard and Pullaiah 2022). We generated whole-genome sequencing data for 55 *Malus* genotypes to at least 15X, as well as 1,216 of their embryos at targets of 3X - 6X depth. These genotypes represent individuals from 22 of the 55 species recognized by Phipps *et al*. and a number of cultivated hybrids, ranging in ploidy from diploid to tetraploid (Phipps et al. 1990; Byrne et al. 2018). We then developed a simple analysis workflow to compare informative biallelic SNPs and detect clonally-produced embryos. We also conducted FCSS on a subset of the sequenced genotypes (N = 26) to compare these data types and understand how they complement one another. Our study is the first to establish low-pass whole genome sequencing as a scalable screening strategy for apomixis. Characterizing reproductive mode is also invaluable for germplasm collection curation, especially when considering secondary and tertiary germplasm for crop breeding. We discuss the advantages and limitations of the method, including ways to make it more efficient and affordable.

## METHODS

### Plant Material

All plant materials (leaves and seed) were obtained from the National Plant Germplasm System (NPGS) *Malus* collection, which is maintained by the United States Department of Agriculture (USDA) at the ARS Plant Genetic Resources Unit in Geneva, New York (National Plant Germplasm System, United States Department of Agriculture-Agricultural Research Service 2025). Accessions were chosen for screening based on a literature review for cases of apomixis in *Malus*. Specifically, we selected individuals of species that had any documentation of producing apomictic seed (e.g., *M. hupehensis, M. sargentii, M. sikkimensis, M. toringo, M. toringoides, M. baccata, M. platycarpa,* and *M. coronaria*), were hybrids with documentation of apomixis (e.g., *M.* ‘Indian Summer’) or had a potential apomictic species in their pedigree (e.g., *M.* ‘Mary Potter’, *M.* ‘Yellow Autumn Crabapple’), were triploid accessions of *M. × domestica* (e.g., cultivars ‘McClintock Grimes’ and ‘Goldgelb 55544’), or were wild diploids with no known apomixis (e.g., *M. floribunda*, and *M. ioensis*). The ploidy level of each maternal genotype was determined previously by NPGS with flow cytometry and is listed with other characteristics of the accession in the GRIN-Global database (Byrne et al. 2018). Information on which accessions were included in the present study, including the number of embryos sequenced and whether there is complimentary FCSS data in the present work, is given in **Supplementary Table 1**.

Open-pollinated seed was isolated from randomly-collected fruits in the fall of 2023 and 2024, then washed in a 30% commercial bleach solution with 0.02% Triton X-100 and rinsed with tap water. It was then stored at 4 °C with drierite until dissection for DNA extraction or for FCSS. Embryos for DNA extraction were dissected using a Motic K-400 Stereo Microscope to ensure its isolation from the seed coat and endosperm. Individual embryos were placed in wells of a 96- deepwell plate (Qiagen, USA, Valencia, CA; cat:19560) on dry ice and stored at -80 °C until DNA extraction. Leaves for DNA isolation of the maternal genotype were either 1) collected in the fall of 2023, lyophilized, and stored dry at room temperature; or 2), were collected fresh in the spring of 2024, flash frozen in liquid nitrogen, and stored at -80 °C until DNA extraction.

### DNA extraction

DNA was extracted directly from embryo tissue to circumvent seed stratification and germination requirements. Up to 96 embryo DNAs were extracted at a time using a modified CTAB protocol (Gasic et al. 2004). Briefly, tissue was ground frozen with 4 mm glass beads using a SPEX SamplePrep 1600 MiniG^®^ (Cole-Parmer sampleprep, Metuchen, NJ), and 500 μL of warm lysis buffer was added directly to the sample and vortexed vigorously. The homogenate was incubated at 55 °C for at least an hour on a plate incubator (Benchmark Scientific Inc., USA, Edison, NJ; model: H6004) set to at least 1000 rpm. Next, 500 μL of 1:1 phenol:chloroform was added, mixed by inversion, then centrifuged at max speed (3486 rcf) in an Eppendorf 5810R bucket centrifuge with attachment A-2-DWP-AT (Eppendorf, Hamburg, Germany). Then 500 μL of a 24:1 chloroform:isoamyl alcohol solution was added to the recovered supernatant and mixed by inversion before another centrifugation step and supernatant recovery. An equal volume of ice-cold isopropanol was added to the recovered supernatant, mixed by inversion, and the DNA was allowed to precipitate overnight in a -20 °C freezer. The following day, DNA was pelleted by centrifugation at max speed for 20 minutes, washed twice with 70-80% ethanol, and eluted in 0.1X TE buffer. DNA quality and concentrations were assessed with a NanoDrop Eight spectrophotometer (ThermoFisher Scientific, Waltham, MA) and a DeNovix model DS-11 FX (DeNovix Inc., Wilmington, DE) with an Invitrogen 1X dsDNA BR kit (ThermoFisher Scientific, Waltham, MA; cat: Q33266), respectively. Most maternal genotype DNAs were extracted in parallel in a separate well alongside their embryos; however, in some cases, maternal DNA was extracted separately using the DNEasy Plant Pro kit (Qiagen USA, Valencia, CA; cat: 69204).

### Library preparation

Libraries for maternal genotypes were prepared as single reactions using either the SeqWell PurePlex High Complexity (Alpha) DNA library prep kit (SeqWell, USA, Beverly, MA; cat: 301400) or the NEBNext® Ultra™ II FS DNA library prep kit for Illumina (New England Biolabs, USA, Ipswich, MA; cat: E7805S) according to the manufacturer’s instructions. DNA inputs ranged between 80ng - 430ng. Libraries for embryos were prepped in pools of up to 96 samples according to the Twist Bioscience 96-plex library prep kit instructions (Twist Bioscience, USA, South San Francisco, CA; part:106543). Each plate was normalized to approximately the same amount of DNA prior to library prep so that DNA inputs ranged between 10ng - 40ng per sample, depending on the plate. Reaction volumes of all reagents were halved, which effectively doubled the amount of samples prepped per kit purchased.

### Sequencing

All samples were sequenced on lanes of a NovaSeq X plus with 10B flow cells in a 150 bp paired-end format at Discovery Life Sciences (Huntsville, AL). Maternal genotypes were pooled in equal concentrations with 16 - 19 individuals per lane. Intended depth for each mother was 28X - 33X assuming a monoploid *Malus* genome size of approximately 650 Mb. For the 6X depth embryo sequencing, batches of up to 96 samples were pooled on one lane each, irrespective of the ploidy levels of each maternal genotype. Once the relationship between average coverage and number of sites was established, the number of samples was increased to 192 samples per lane, with an intended depth of about 3X per embryo. Pooling concentrations for the 3X depth sequencing was intentionally favored toward libraries with more polyploids, in proportion to the total number of maternal haplotypes per plate. This was to ensure potentially higher ploidy embryos would receive more sequencing for calling for better accuracy. Plates of embryos were demultiplexed according to their i7 adapter sequence after sequencing, and fgbio version 2.2.1 was used to demultiplex individual samples on a plate based on their i5 sequence (Fennell and Homer 2024). All sequencing data is available for download on NCBI’s SRA under BioProject number (XXXXXX).

### Variant calling pipeline

Fastp version 0.23.4 was used to trim and remove adapters, remove reads with too many low-quality bases (>30%), and deduplicate the demultiplexed, compressed raw fastq files (Chen et al. 2018). Reads for each sample were aligned to haplome A of the *Malus × domestica* ‘Honeycrisp’ v1.1.a1 genome with bwa-mem version 0.7.17 (Li and Durbin 2009). The genome and annotation was downloaded from <u>rosaceae.org</u> (Jung et al. 2019). Samtools version 1.19.2 was used to extract raw and filtered read alignments and calculate average depth, as well as sort and filter the alignments (Danecek et al. 2021). Average depth was estimated for each sample by multiplying the number of aligned reads by 150 (the maximum read length) and dividing it by the monoploid apple genome size in base pairs (650,000,000). Only alignments with mapQ greater than 30 were used for variant calling, and secondary alignments (multi-mapping reads) were excluded.

Both bcftools (v1.19) and freebayes (v1.3.1) were used to call genotypes to enhance accuracy (Garrison and Marth 2012; Danecek et al. 2021). Each sample was called and filtered separately, and only biallelic SNPs were considered. To assess the accuracy of the genotype calls in the 6X depth experiments, we subsetted the maternal genotype’s trimmed fastq file at 2X, 4X, and 12X depth to create “simulated” clonal embryos in the ranges of the actual embryos’ depths. For the 3X experiments, we created simulated embryos at depths of 1X and 3X. We optimized the noise-to-signal ratio by iteratively testing different variant quality filters, noting the number of mismatches between simulated embryos and the full maternal dataset, exploring the reasons for mismatches through manual examination in IGV version 2.13.0 (Thorvaldsdóttir et al. 2013), and adjusting the filters accordingly. It became evident that the simulated embryos with lower depths were more sensitive to noise, as reflected by a slightly higher error rate in the variant calls (max error rate = 3.6 %; adjusted r^2^ = 0.54 between simulated embryo depth and error). On the other hand, sites with abnormally high coverage in low-depth libraries could reflect problems in alignments. Low (LDP) and high depth (HDP) filters, or the minimum and maximum number of reads supporting a site, were set according to the ploidy of the mother and the embryo’s approximate depth. For embryos of diploid, triploid, and tetraploid mothers, a minimum of 10, 11, and 12 reads, respectively, were required to support a genotype call. For embryos with approximate depths of 1X, 2X, 3X - 6X, and greater than 6X, the maximum number of reads allowed for any given call was 25, 30, 30, and 40, respectively. Maternal genotypes’ LDP cutoff was set to 15, and the HDP was set to five times the filtered depth. In addition to the depth filters, only variants with genotype qualities (GQ) greater than 30 and site qualities (QUAL) greater than 50 were kept for downstream analysis.

After calling and filtering, we used the SelectVariants –concordance function from GATK version 4.6.0.0 to keep sites that were present in both the bcftools and freebayes vcf files (McKenna et al. 2010). Concordant embryo files were pruned with bcftools +prune, selecting only one site per 1000 bp to reduce bias in the results due to linkage. A limitation of the pruning function allowed only the preservation of variant sites relative to the reference genome (0/1 or 1/1 locations); our controls demonstrated this did not affect our ability to detect clonal embryos. Afterwards, the concordant, filtered, maternal vcf was merged with her concordant pruned embryo vcfs. Bcftools gtcheck was used to determine the number of mismatching sites between mother and every child. Gtcheck automatically skips over potentially uninformative sites, where all individuals in the merged vcf are either homozygous reference (0/0) or missing (./.).

### Visualization

Data were imported into R version 4.2.1, and tidyverse, ggplot2, lme4 packages were used for data reformatting, generating plots, and performing statistical analyses (Bates et al. 2015; Wickham 2016; Wickham et al. 2019; R Core Team 2021). The association of maternal genotype and depth was only examined for samples sequenced to approximately 6X depth, as the lower depth sequencing (3X) was intentionally skewed to give embryos derived from polyploid mothers more sequencing resources per lane. Sequencing results are summarized in **Supplementary Table 2**, and all scripts are available at https://github.com/goeckeritz/high-throughput-sequencing-screen-for-apomixis.

### Nuclei isolation and preparation for FCSS

668 seeds of 12 *Malus* species and several cultivated hybrids (represented by 26 individuals; **Supplementary Table 3**) were first prepared from pomes and cut in half. Only well-developed seeds were used for the detection of reproductive modes based on a FCSS (Matzk et al. 2000). Each sample consisted of one half of seed with the internal standard (*Carex acutiformis* 2C = 0.82 pg; (Temsch et al. 2021)). DAPI (4’,6-diamidino-2-phenylindole) staining was used with a Partec CyFlow ML cytometer equipped with a 365-nm UV LED. A simplified two-step protocol was followed (Dolezel et al. 2007). A slight modification consisting of using a larger amount of the Otto I buffer (0.7 ml; according to (Macková et al. 2020)) and shortening of incubation time (max 2 min) was adopted. The ploidy of the embryo and the endosperm was calculated from the peaks of the fluorescence histograms.

## RESULTS

### Sequencing statistics and adjusting depth of sequencing

To develop a low-pass WGS screen for apomixis, *Malus* embryos from 16 maternal genotypes were first sequenced at an average depth of 6X with up to 96 samples per NovaSeq X Plus 10B lane (termed the 6X embryos; n = 355). The distribution of average read depth for these embryos is shown in **Figure 1A**. Depth ranged widely from 0.3X to 23.3X, and the mean and median a was 6.1 and 5.9, respectively, with several samples at average read depths greater than 2.5 standard deviations above the mean driving the right skew. A linear regression of average depth to the maternal genotype showed a statistically significant association between these two variables (p=1.907x10^-7^, type III ANOVA), suggesting the sequencing performance of each embryo was dependent on the maternal genotype it came from (**Supplementary Figure 1**).

**Figure 1:**
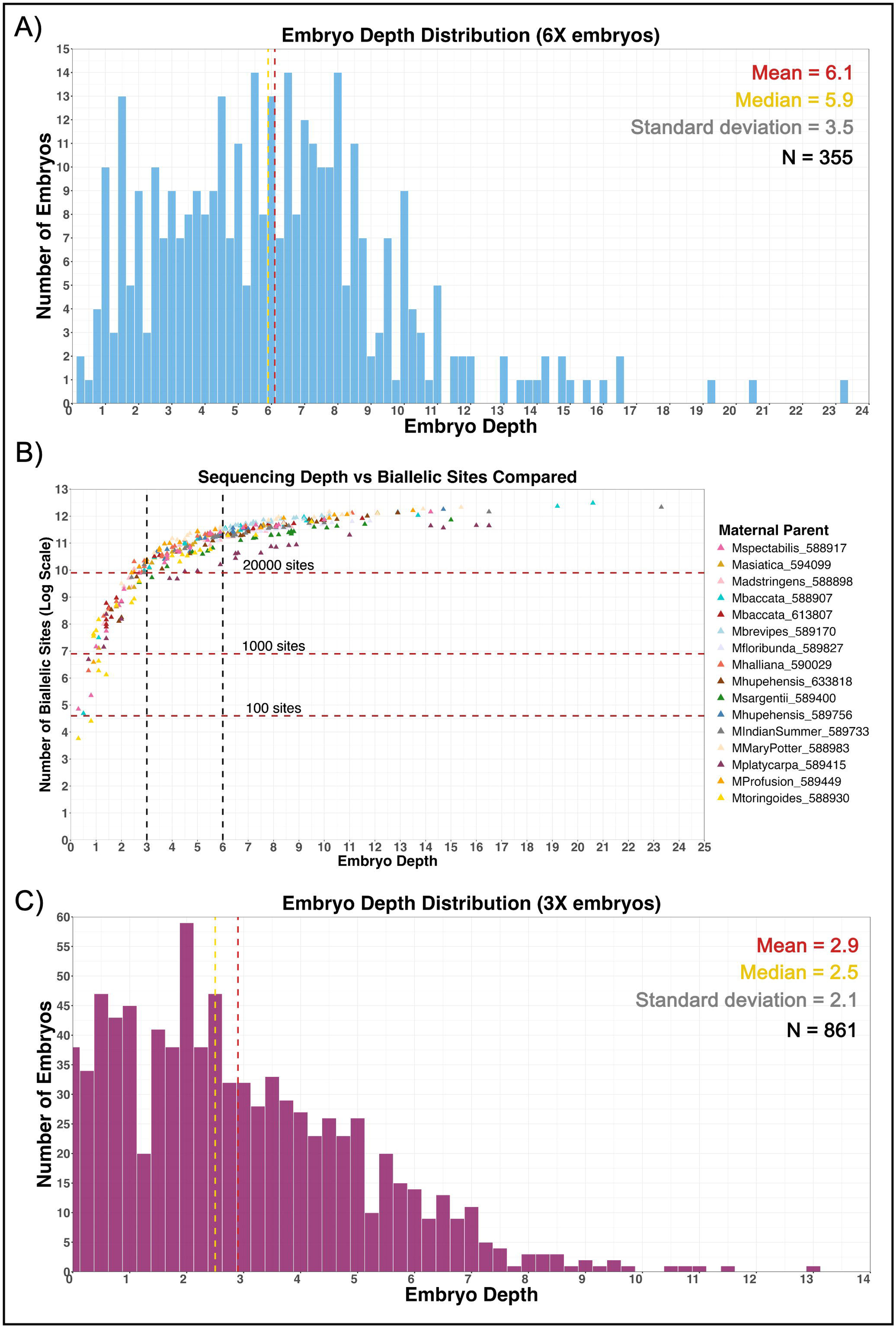
**A)** Distribution of average depths for all embryos that were sequenced at a target of 6X depth (up to 96 samples per lane). Average depth was calculated as (# of fastp-processed reads aligned to the reference genome)*150 / 650000000), or the number of trimmed aligned reads multiplied by the read length format, then divided by a typical monoploid genome size for *Malus*. Bin size = 0.25. **B)** Relationship between average raw sequencing depth and the number of high-quality, biallelic variant sites compared between an embryo and its mother (natural log scale). The target depths of the present study are marked (3X, 6X) in addition to y-values corresponding to 100, 1000, and 20,000 sites compared. **C)** Distribution of average depths for all embryos that were sequenced at an average of 3X depth. Average depth for every embryo was calculated similarly to those sequenced at 6X target depth. Bin size = 0.25.

We used the 6X sequencing data to depict the relationship between the number of biallelic sites compared (after quality filtering alignments, calls, and pruning) and average raw sequencing depth to decide whether sequencing at lower depths would be a cost-efficient alternative (**Figure 1B**). Based on this association, we reasoned embryos sequenced at an average of 3X would still provide hundreds to thousands of informative sites to determine clonality. Notably, even lower depths (e.g., 1.5X) were estimated to provide over 1000 sites for comparison between mother and child; however, given the high variability in sequencing depth between embryos and to minimize sample drop-out, we moved forward with average sequencing depths of 3X per embryo. The compared sites were randomly distributed along the lengths of the 17 *Malus* reference chromosomes (**Supplementary Figure 2**).

A total of 40 genotypes’ embryos were sequenced at an average of 3X depth (termed the 3X embryos; n=861), with actual depths ranging between 0X to 13.1X (**Figure 1C; Supplementary Table 1**). *M. platycarpa* 589415 was screened twice given the discrepancies first seen in the 2023 sequencing and 2024 FCSS data, bringing the total number of screened genotypes to 55 and the number of embryos sequenced to 1,216. The mean, median, and standard deviation for depth of 3X embryos were 2.9, 2.5, and 2.1, respectively. Two 6X embryos and 58 3X embryos were excluded from analysis as fewer than 100 high-quality sites could be compared to their maternal library.

### Accounting for sources of error

We used a combination of approaches to account for sources of error in the sequencing data. First, we included known obligate sexually-reproducing genotypes (*M. spectabilis* 588917 and *M. asiatica* 594099) and facultatively apomictic genotypes with high penetrance (*M. hupehensis* 633818 and *M. sargentii* 589400) in the 6X depth sequencing experiments. Aposporic apomixis in *Malus* is facultative, so even highly-penetrant apomicts are expected to produce some proportion of non-clonal embryos. High-penetrant apomicts are defined here as producing > 50% clonal progeny. These controls allowed us to differentiate true clonal embryos in the presence of most other sources of noise. Second, for most genotypes (n = 40), one to three maternal DNA extracts were processed in parallel with her embryo DNAs; these samples are referred to as ‘spike-ins.’ Spike-ins allowed us to document any error that may have risen from the differences in sequencing library preparation approach between mother and child. Finally, all 55 maternal libraries were subsampled at several depths representative of the population of embryos used to screen them for reproductive mode. These subsets, or ‘simulated embryos’, provided an estimate of variant calling error at different depths when compared to the full maternal library dataset.

#### 1) Reproductive mode controls

The screening results for four control genotypes with known types of reproduction (obligate sex or facultative apomixis) are shown in **Figure 2**, alongside representative flow cytometry spectra for each. The genetic differences between the maternal genotype and each of her embryos was visualized as a percent of matching sites where she was called homozygous (0/0 or 1/1; y-axis in **Figure 2A**), and the percent matching sites where the maternal genotype was called heterozygous (0/1; x-axis). Thus we could quickly visualize relative similarities after the addition of new alleles (outcrossing) and recombination, respectively. As expected, the sexual controls (*M. spectabilis* 588917 and *M. asiatica* 594099) produced embryos with dispersed similarities compared to the maternal genotype while facultative apomixis controls (*M. hupehensis* 633818 and *M. sargentii* 589400) showed a high proportion of embryos clustering with nearly 100 % matches on both axes (**Figure 2A**, **Supplementary Table 2**). These results were consistent with the flow cytometry seed screen (FCSS) (**Figure 2B**, **Table 1**, **Supplementary Figure 3**, **Supplementary Table 3**). All seed analyzed by FCSS in 2023 and 2024 for the sexual controls were indicative of sexual reproduction, with the exception of one seed from *M. asiatica* 594099. In contrast, seed analyzed for the facultative apomixis controls indicated a mix of seed produced through ‘pure’ apomixis (apospory + parthenogenesis), B_III_ hybridization (genome increase through pollination of an unreduced egg) and sex (gamete reduction + pollination).

**Figure 2:**
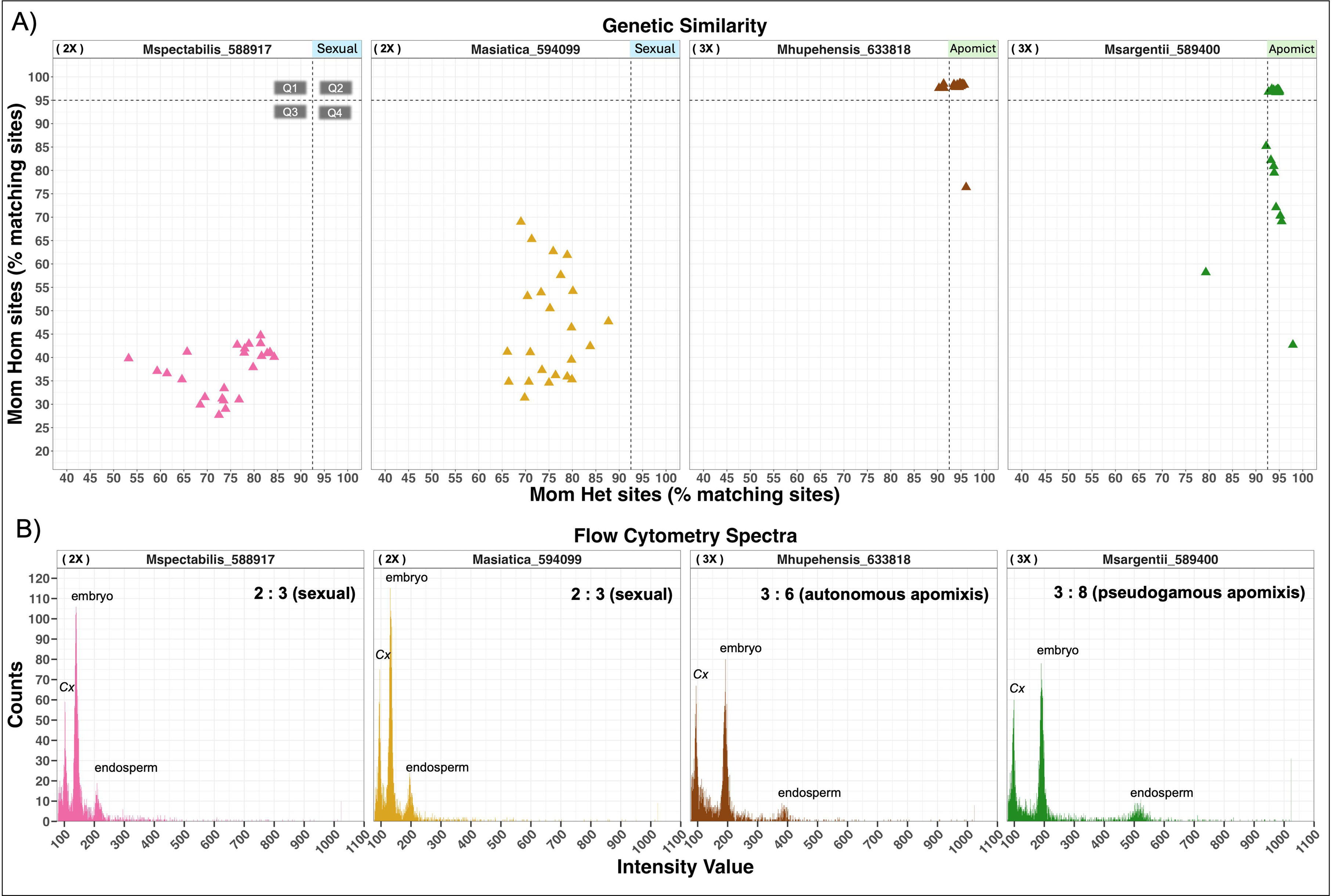
**A)** Screening results for genotypes known to reproduce via sex (*M. spectabilis* 588917 and *M. asiatica* 594099) and apomixis (*M. hupehensis* 633818 and *M. sargentii* 589400). Each colored triangle represents a single embryo sequenced to 6X depth. Percent similarity of the embryo compared to the maternal genotype is plotted according to the type of call. Similarity between embryo and mother when she was called as homozygous (0/0 or 1/1) for a biallelic site is reflected on the y-axis, while similarity between embryo and mother when she was called as heterozygous (0/1) is shown on the x-axis. Black dashed lines (x=92.5%, y=95%) delineate Q1, Q2, Q3, and Q4 and the reproductive scenarios each represents are described in the text. A genotype was required to produce at least one clonal embryo (Q2) to be a facultative apomict; otherwise, the genotype was presumed to reproduce via sexual processes. Included above each subplot is the ploidy level and the reproductive mode on the left and right, respectively, of the genotype’s ID. **B)** A representative flow cytometry spectrum from each genotype showing the predominant mode of reproduction. The internal standard (*Cx*), embryo, and endosperm peaks are labeled in each subplot. The approximate ratio of embryo to endosperm ploidy is given with the predicted mode of reproduction for that seed near the top right.

**Table 1:**
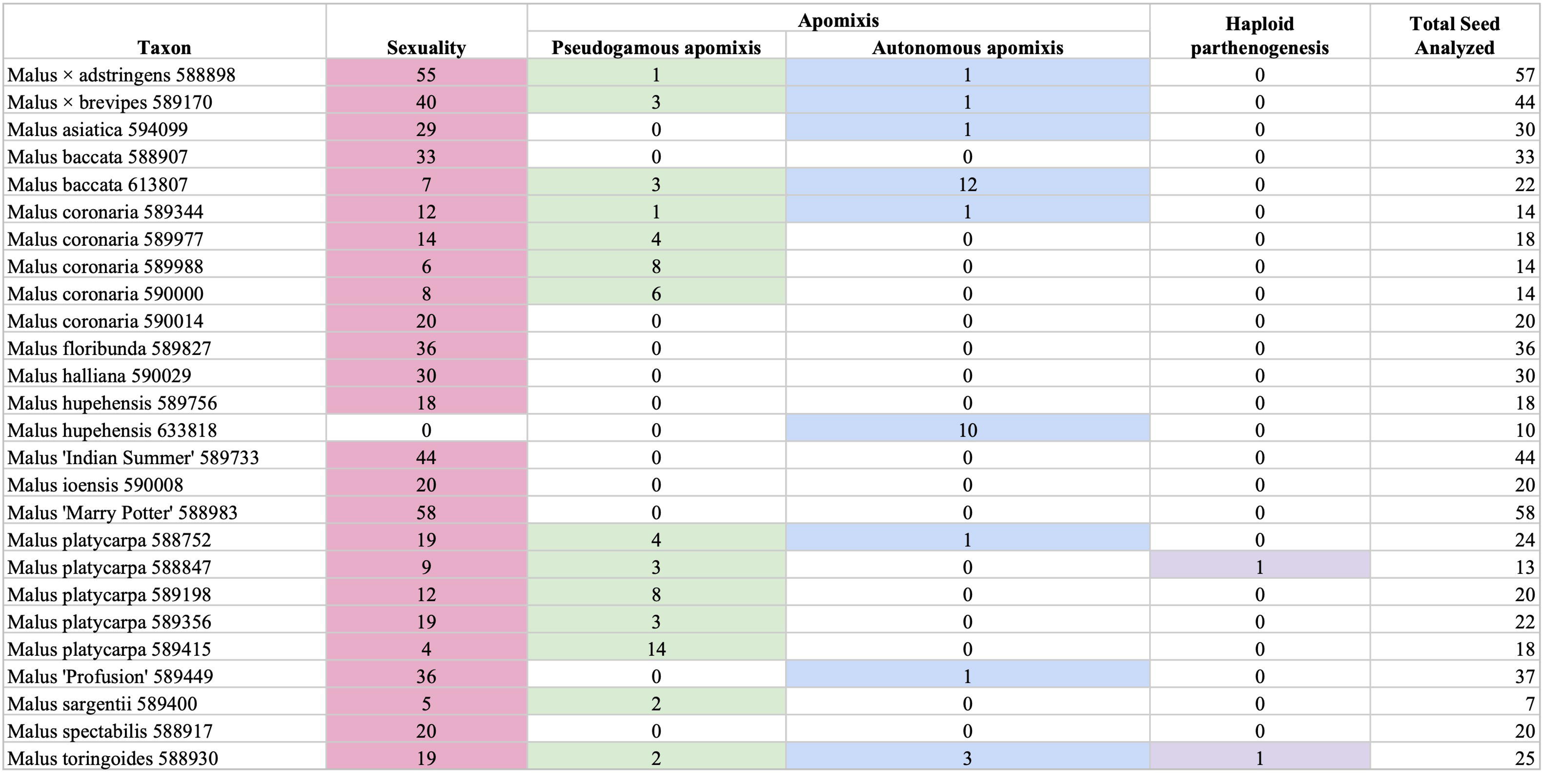
Flow cytometry seed screen (FCSS) summary for 26 apple genotypes. **Taxon** - Genotype and PI number in the USDA germplasm database; **Sexuality** - seed counts exhibiting embryo:endosperm (Emb:End) ratios reflective of syngamy / sexual reproduction. This includes reduction + fertilization or genome increase scenarios, where unreduced eggs are fertilized with self or foreign pollen (B_III_ hybrids). **Pseudogamous apomixis** - seed counts exhibiting Emb:End ratios reflective of pseudogamous apomixis, where ploidy of the embryo is the same as the mother and endosperm = (2*Emb + Paternal contribution). **Autonomous apomixis** - seed counts exhibiting Emb:End ratios reflective of autonomous apomixis, where ploidy of the embryo is the same as the mother and the endosperm = (2*Emb). **Haploid parthenogenesis** - seed counts exhibiting Emb:End ratios reflective of pseudogamous apomixis, where ploidy of the embryo is half the mother’s and the endosperm = (2*Emb + Paternal contribution). **Total Seed Analyzed** - total seeds analyzed by FCSS in both years (2023 and 2024). See **Supplementary** Table 3 for more information.

| Taxon | Sexuality | Apomixis |  | Haploid<br>parthenogenesis | Total Seed<br>Analyzed |
| --- | --- | --- | --- | --- | --- |
|  |  | Pseudogamous apomixis | Autonomous apomixis |  |  |
| Malus × adstringens 588898 | 55 | 1 | 1 | 0 | 57 |
| Malus × brevipes 589170 | 40 | 3 | 1 | 0 | 44 |
| Malus asiatica 594099 | 29 | 0 | 1 | 0 | 30 |
| Malus baccata 588907 | 33 | 0 | 0 | 0 | 33 |
| Malus baccata 613807 | 7 | 3 | 12 | 0 | 22 |
| Malus coronaria 589344 | 12 | 1 | 1 | 0 | 14 |
| Malus coronaria 589977 | 14 | 4 | 0 | 0 | 18 |
| Malus coronaria 589988 | 6 | 8 | 0 | 0 | 14 |
| Malus coronaria 590000 | 8 | 6 | 0 | 0 | 14 |
| Malus coronaria 590014 | 20 | 0 | 0 | 0 | 20 |
| Malus floribunda 589827 | 36 | 0 | 0 | 0 | 36 |
| Malus halliana 590029 | 30 | 0 | 0 | 0 | 30 |
| Malus hupehensis 589756 | 18 | 0 | 0 | 0 | 18 |
| Malus hupehensis 633818 | 0 | 0 | 10 | 0 | 10 |
| Malus 'Indian Summer' 589733 | 44 | 0 | 0 | 0 | 44 |
| Malus ioensis 590008 | 20 | 0 | 0 | 0 | 20 |
| Malus 'Marry Potter' 588983 | 58 | 0 | 0 | 0 | 58 |
| Malus platycarpa 588752 | 19 | 4 | 1 | 0 | 24 |
| Malus platycarpa 588847 | 9 | 3 | 0 | 1 | 13 |
| Malus platycarpa 589198 | 12 | 8 | 0 | 0 | 20 |
| Malus platycarpa 589356 | 19 | 3 | 0 | 0 | 22 |
| Malus platycarpa 589415 | 4 | 14 | 0 | 0 | 18 |
| Malus 'Profusion' 589449 | 36 | 0 | 1 | 0 | 37 |
| Malus sargentii 589400 | 5 | 2 | 0 | 0 | 7 |
| Malus spectabilis 588917 | 20 | 0 | 0 | 0 | 20 |
| Malus toringoides 588930 | 19 | 2 | 3 | 1 | 25 |

We next examined the ranges in percent similarities for clonal embryos from our facultatively apomictic controls. In *M. hupehensis* 633818, one embryo obviously deviating from the main cluster on the y-axis and three slightly deviating on the x-axis were excluded from these range considerations. In *M. sargentii* 589400, nine deviating from the tight cluster at nearly 100 % similarities were excluded from range considerations. After these conservative exclusions, similarities for 33 clonal embryos ranged between 94.6 % - 96.4 %. Notably, percent similarities for heteroallelic sites (0/1) trended several points lower than for homoallelic sites (0/0 or 1/1), likely due to the requirement that both alleles were sampled at appropriate depths for correct calling. The range of heteroallelic site similarities for true clonal embryos was 92.7 % - 95.7 % and for 96.7 % - 98.6 % for homoallelic site similarities. A 95 % confidence interval based on quantiles for the 33 embryos placed the heteroallelic lower boundary at 93.3 % and the homoallelic lower boundary at 96.7 %.

#### 2) Spike-ins

To estimate the potential error rate associated with library prep between mother (SeqWell or NEBNext; **Supplementary Table 1**) and embryo (Twist Biosciences 96-plex), one to three replicates of the mother’s leaf DNA extract was prepped in parallel in the same batch with her embryos. In total, we analyzed spike-in information from 38 maternal genotypes after variant filtering, representing a range of ploidies and *Malus* species. A histogram of percent similarities for 85 total spike-ins with at least 100 sites compared revealed a clear bimodal distribution (**Supplementary Figure 4**). Most spike-ins showed > 93 % similarity to their deeper sequenced library prepared with SeqWell or NEBNext, therefore providing confidence in comparing library types (SeqWell or NEBNext versus 96-plex Twist Bioscience); however, eight showed similarities ≤ 88.1 %. Upon closer examination, all eight samples belonged to the four tetraploid *M. platycarpa* accessions screened in the present study. The FCSS data indicated all four of these individuals were facultative apomicts, or capable of producing clonal embryos (**Table 1, Supplementary Figure 3, Supplementary Table 3**). To rule out library preparation bias (between mother and child) being greater only in these accessions, we evaluated one pair of spike-ins for each of the four *M. platycarpa* accessions against one another, so that the same leaf DNA extracts prepared with Twist Bioscience were compared to one another. At the same time, we compared a pair of spike-ins from three other tetraploids: *M. coronaria* 589344 *, M. sikkimensis* 589599, and *M. sargentii* 589372. These spike-ins were sequenced to adequate depths (> 3.7X) for same-library comparisons.

Despite being prepared from the same DNA extract and same library kit, *M. platycarpa* spike-in comparison similarities still lagged 7% - 8% lower than the other tetraploid genotypes. Thus, while some library preparation bias may be present in samples, the larger genetic discrepancies between the *M. platycarpa* spike-ins and their deeper-sequenced libraries prepared with SeqWell or NEBNext kits likely have an unknown biological explanation. The tetraploid *M. platycarpa* spike-in range of similarities for heteroallelic sites was 86.4 % - 88.9 % and 82.1 % - 84.7 % for homoallelic sites. For all other samples, the heteroallelic and homoallelic ranges were 91.4 % -98.4 % and 91.4 % - 99.6 %, respectively.

#### 3) Simulated embryos

Subsampled depths for simulated embryos (n = 321 after filtering for at least 100 sites compared) varied from 1.2X - 22.5X, and the range for heteroallelic similarities in simulated embryos was 95.4% - 100% and was 98.5% - 100% for homoallelic similarities. Although there was greater variability in simulated embryos subsampled at lower depths, the > 95 % similarities between the subsampled reads and the entire maternal library indicated a high call accuracy. Still, it should be noted that in some cases the variant calling was erroneous even as the same read data was sampled.

### Estimating boundaries for clonal embryos

Taking these sources of error into account, we determined lower cutoffs for heteroallelic (92.5 %) and homoallelic (95 %) sites when determining clonal embryos from other reproductive scenarios. For tetraploid *M. platycarpa* accessions, boundaries were readjusted to 85 % for heteroallelic sites and 82 % for homoallelic sites. The four quadrants delineated by these cutoffs (**Figure 2A**; *M. spectabilis* 588917 subplot) were predicted to represent the following types of reproduction: gamete reduction, recombination, and self-pollination or autogamy (Q1), apomixis, including apomeiosis and parthenogenesis (Q2), gamete reduction, recombination, and foreign pollination or xenogamy (Q3), and either gamete reduction and fixed haplotypes/low recombination among preferred mating partners or the foreign pollination of an apomeiotic egg cell (Q4). Notably, there was also the possibility that embryos produced by haploid parthenogenesis (e.g., AABB → BB and AAAA → AA) could appear in Q1, and that self-pollinated B_III_ embryos (self-pollinated apomeiotic embryo sacs; e.g., AAB → AABB and AAA→ AAAA) could aggregate in Q2 as well. However, only two haploid parthenogenesis events were captured with FCSS, thus this type of reproduction may be rare in *Malus* (**Table 1**, **Supplementary Table 3, Supplementary Figure 3**). In the case of self-pollinated B_III_ embryos, FCSS could capture ploidy increases but would not be able to differentiate self or outcrossed pollination. Nevertheless, since individuals with putative B_III_ embryos (distinguished by FCSS) commonly displayed some level of apomixis, we speculated self pollinated B_III_ embryos were unlikely to lead to false positives in identifying facultative apomicts. A genotype was determined to reproduce with facultative apomixis via the sequencing screen if at least one embryo derived from that genotype appeared in Q2.

### Screening-by-sequencing and FCSS results

The screening-by-sequencing results for 51 *Malus* genotypes maintained by the USDA in Geneva, NY are shown in **Figure 3** (also see **Supplementary Table 2**). The majority of these genotypes (n = 29) only produced embryos through putative sexual processes (Q1, Q3, and Q4). In accordance with *Malus* species mainly reproducing through xenogamy (obligate outcrossing), the majority of embryos from these genotypes fell in Q3. Still, rare selfing events do occur in *Malus* and polyploidy can lead to self-compatibility (Dickinson et al. 2007). Concordant with these facts, a few selfing events (Q1) were detected in diploids (*M. baccata* 588907, *M.* ‘Robinson’ 589455, *M.* ‘Ralph Shay’ 589734), but most autogamy was detected in triploids and tetraploids. The flow cytometry data from obligate sexual individuals largely supported the sequencing results (embryo : endosperm DNA ratios were approximately 2 : 3), but also detected rare instances of potential apomixis or unreduced gamete formation (e.g., in *M. asiatica* 594099, *M.* ‘Profusion’ 589449, *M. brevipes* 589170, and *M. × adstringens* 588898) and genome increases from possible mixed-ploidy crosses (e.g., *M. baccata* 588907; **Table 1**, **Supplementary Figure 3**, **Supplementary Table 3**). All obligate sexually-reproducing individuals are diploids except for *M. × domestica ‘*Goldgelb 55544’ 589458 and *M. × domestica* ‘McClintock Grimes’ 589124, which are both triploids. These findings are consistent with the notion that cultivated apples, irrespective of ploidy, reproduce through sexual processes (Bergström 1938; Howard et al. 2022).

**Figure 3:**
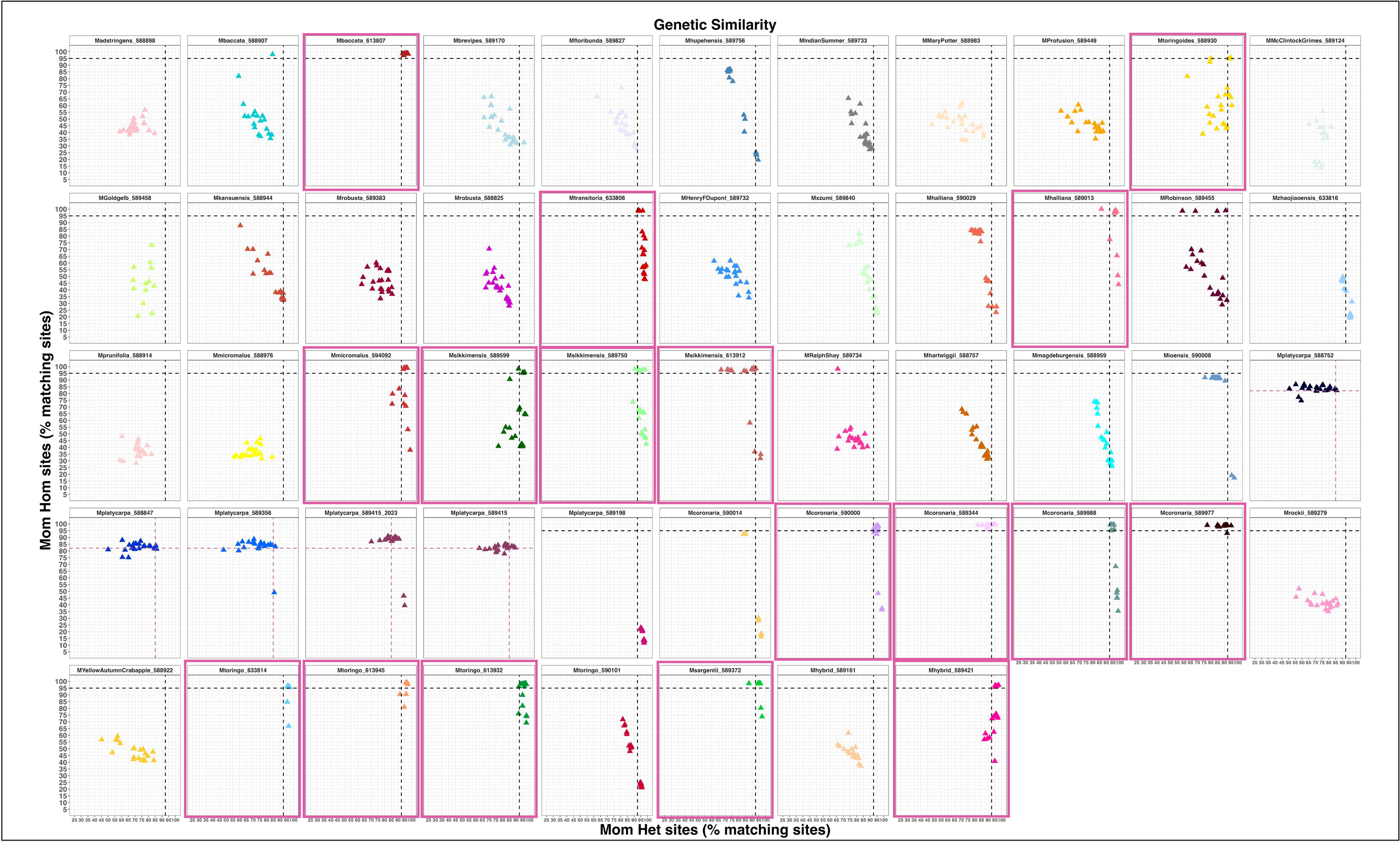
Screening-by-sequencing results for 51 *Malus* genotypes maintained by the USDA in Geneva, NY. Each colored triangle represents a single embryo sequenced to a target of 3X – 6X depth. Percent similarity of the embryo compared to the maternal genotype is plotted according to the type of call. Similarity between embryo and mother when she was called as homozygous (0/0 or 1/1) for a biallelic site is reflected on the y-axis, while similarity between embryo and mother when she was called as heterozygous (0/1) is shown on the x-axis. Black dashed lines (x=92.5%, y=95%) and pink dashed lines (x = 85%, y = 82%; tetraploid *M. platycarpa* accessions only) within each plot delineate Q1, Q2, Q3, and Q4 and the reproductive scenarios they represent are described in the text. A genotype was required to produce at least one clonal embryo (Q2) to be a facultative apomict; otherwise, the genotype was presumed to reproduce via sexual processes. Plots boxed in pink highlight facultatively apomictic genotypes. Plots boxed in blue highlight data from tetraploid *M. platycarpa* genotypes, which were confirmed to be facultative apomicts with FCSS. *Malus platycarpa* 589198 is boxed in gold to point out this genotype produced embryos only in Q4 in 2023, but FCSS data from 2024 suggested it reproduces by both apomixis and B_III_ hybridization. Note *Malus platycarpa* 589415 embryos were sequenced in both 2023 and 2024. Ploidy, number of embryos sequenced, and whether a genotype has complementary FCSS data is reported in **Supplementary Table 1**.

The skim-seq screen identified 17 facultative apomicts, 8 of which were complemented with FCSS and shown to produce clonal embryos via apomixis (autonomous and/or pseudogamous; **Figure 3**, **Table 1**). Disagreements between the sequencing data (prior to readjusting quadrant cutoffs) and FCSS belonged to five *M. platycarpa* accessions; the four tetraploid individuals (blue boxes in **Figure 3**) consistently showed large differences even between internal sequencing controls as discussed above. However, triploid *M. platycarpa* 589198 only produced embryos in Q4 according to those sequenced in 2023, while FCSS results from 2024 indicated both pseudogamous apomixis and genome increase (fertilization of an unreduced embryo sac) as its prospective reproductive routes. As *M. platycarpa* 589198 did not show the same spike-in trends as tetraploid genotypes, we propose the absence of clonal embryos in the sequencing screen of 2023 embryos is not due to a technical limitation of the method. Rather, we hypothesize spring conditions differing between years may have influenced the reproductive balance between pure apomixis and genome increase in this individual given that apomixis is known to be influenced by environmental factors such as temperature (Liu et al. 2014; Klatt et al. 2018).

All facultative apomicts screened here are either triploid or tetraploid except for *M.* hybrid (*M. rockii*) 589421, which is diploid and documented to be phenotypically similar to *M. sikkimensis* in the grin-global database (Byrne et al. 2018). Most apomictic accessions belong to species native to eastern Asia and North America that have previously been documented to exhibit apomixis among their populations: *M. hupehensis*, *M. sargentii*, *M. baccata*, *M. sikkimensis*, *M. toringoides*, *M. transitoria*, *M. platycarpa*, and *M. coronaria* (Sax 1959; Kron and Husband 2009). However, species classifications did not perfectly correspond to reproductive mode. For example, *M. baccata* 613807 reproduced through facultative apomixis according to both the sequencing screen and FCSS results while *M. baccata* 588907 was shown to only produce sexual embryos (**Figure 3**).

Interestingly, triploid apomicts *M. sargentii* 589400 (**Figure 2A**), *M. transitoria* 633806, *M. halliana* 589013, *M. micromalus* 594092, *M. sikkimensis* 589750, *M. platycarpa* 589198, *M. coronaria* 590000, *M. coronaria* 589988, *M. toringo* 633814, *M. toringo* 613945, and *M. toringo* 613942 mainly produced embryos in Q2 or Q4 (**Figure 3**). In addition to *M. platycarpa* 598198, available FCSS data from *M. sargentii* 589400, *M. coronaria* 590000, and *M. coronaria* 589988 confirmed pseudogamous apomixis and genome increase as the predominant modes of reproduction, strongly implying Q4 may represent apomeiosis and foreign pollination (B_III_ hybridization). Still, complementary FCSS observations from *M. hupehensis* 589756 and *M. halliana* 590029, which also produced embryos in Q4, showed these individuals to reproduce sexually and therefore Q4 also could represent fixed alleles/haplotype blocks or lower rates of recombination among mating partners as postulated above. In contrast to triploid apomicts, tetraploid apomicts (*M. toringoides* 588930, *M. sikkimensis* 589599, *M. sikkimensis* 613912, *M. coronaria* 589344, *M. coronaria* 589977, *M. sargentii* 589372, and tetraploid *M. platycarpa* individuals) regularly produced selfed embryos (Q1), which may be an indirect consequence of pollen sterility in triploid apples.

## DISCUSSION

In the present study, we took advantage of the immense genetic resources maintained by the USDA to develop a skim-sequencing approach and detect apomixis (clonal seed production) in *Malus* genotypes. A total of 55 individuals representing over 20 recognized species, hybrids, and cultivated apples were screened by sequencing embryos at low depth and comparing mother and progeny sequence data. A subset of these individuals (N = 26) was also analyzed by FCSS to validate and complement the skim-seq approach, and with the exception of four genotypes before quadrant readjustment, these data types were consistent with one another. While hosts of studies have used molecular markers to detect clonal reproduction in populations, our study is the first to illustrate how high-throughput, WGS skim-sequencing approaches are a viable option to screen for apomixis.

We used plate-based DNA extractions and Twist Bioscience’s 96-plex library kit to create libraries for over 1,200 open-pollinated embryos. *Malus* genomes are relatively small (650 Mb), so the combined costs of library preparation (using half-volume reactions) and sequencing each embryo to 3X depth came to roughly $10.50 per sample. The relationship between the number of sites analyzed and sequencing depth indicates that reliable conclusions could also be drawn at 1.5X depth (**Figure 1B**), although excellent sample normalization would be required to minimize substantial sample dropout. Ploidy, allele sampling probabilities, and heterozygosity levels should also be considered when designing these experiments. While still relatively expensive for labs that may be well-equipped to conduct flow cytometry, we expect declining costs, improvements in sequencing chemistries, and DNA extraction automation to lead to these approaches becoming more commonplace for investigating reproductive plant biology.

The flow cytometry seed screen (FCSS) was developed 25 years ago and remains the dominant methodology to detect apomixis and other modes of reproduction in plants (Matzk et al. 2000). It takes advantage of the process of double fertilization and relative DNA quantities between embryo and endosperm to indirectly determine reproductive types. Sequencing offers a more direct approach to identifying clonal and sexual progeny but has been prohibitively expensive until recently. We also demonstrated the importance of using internal sequencing controls when determining clonal embryo cutoffs for some genotypes (*M. platycarpa* tetraploids). While both FCSS and sequencing methods have their advantages and disadvantages, together they yield an exceptionally comprehensive view into plant reproductive biology. For example, FCSS may distinguish between autonomous and pseudogamous forms of apomixis and can clearly detect ploidy changes in progeny. Conversely, sequencing-based approaches can easily discern self-fertilization (autogamy) from outcrossing events (xenogamy). Together, these approaches allowed us to create more complete breeding profiles for genotypes than either would have alone.

Interestingly, we observed rare (10% or less of FCSS seed analyzed for a genotype) apomixis-like events in certain diploid individuals, including *M. × adstringens* 588898, *M. brevipes* 589170, *M.* ‘Profusion’ 589449, and *M. asiatica* 594099, yet all embryos examined with the sequencing screen showed genetics reflective of gamete reduction and/or recombination and cross-pollination (Q3, one embryo in Q4 from *M. brevipes*; **Table 1, Supplementary Figure 3,** **Figure 3**). While this could simply be a result of sampling, many plant species including *Malus* are known to produce occasional unreduced gametes through meiotic abnormalities such as first or second division restitution (Bretagnolle and Thompson 1995; Howard et al. 2022). Applying FCSS and genotyping methods to the same seed would provide information on whether these rare cases were truly clonal (apomictic) or formed by some other means, as has been done for *Rubus* (Šarhanová et al. 2024). Classical developmental staging using confocal microscopy and advanced imaging technologies would be informative on the mechanism, as was elegantly shown in several Asteraceae taxa (Cornaro et al. 2025). In summary, emerging multi faceted approaches promise to resolve the environmental, genetic, and epigenetic influences on plant reproduction with unmatched precision.

The distributions of embryos’ genetic similarities to their maternal parent in **Figure 3** highlights questions regarding apple mating systems, including those on pollen competition and interspecific barriers. For instance, the detection of rare selfing events in *Malus* implies that even if individuals are self-compatible, xenogamy typically prevails over autogamy. Although this phenomenon has been widely documented, the underlying genetic and environmental causes are complex and vary between species (Whitehead et al. 2018). Our experimental conditions included *Malus* species from around the world planted together in a common orchard, which increased the likelihood of mating among individuals that would not normally encounter one another in nature. Yet, certain North American sexuals (*M. ioensis* 590008, *M. coronaria* 590014) and facultative apomicts (polyploid *M. platycarpa* and *M. coronaria* individuals) show highly constrained distributions that may imply preferential mating. *Malus coronaria* and *M. platycarpa* in particular exhibit discordant phylogenetic signals and have been described as cryptic or recent hybrids of divergent *Malus* lineages (Liu et al. 2022; Li et al. 2025; Zhang et al. 2025). Perhaps reflective of their putative hybrid genome structure, tetraploid North American species (*M. platycarpa* in blue boxes of **Fig. 3**, *M. coronaria* 589344, and *M. coronaria* 589977) deviate mainly on the x-axis, indicating apomixis and selfing to be their primary modes of reproduction despite the abundance of intra- and interspecies partners. Better representation of these species in phylogenetic studies is sorely needed to elucidate their evolutionary relationships within the greater *Malus* clade.

*Malus* is a taxonomically challenging genus of over 50 named species with reticulate evolutionary processes such as hybridization and polyploidy in addition to apomixis (Phipps et al. 1990; Sun et al. 2020). Gametophytic apomixis in *Malus* is dominant and the genetic components regulating apomeiosis and parthenogenesis are thought to act separately, which has also been demonstrated in dandelion and *Cenchrus* (Sax 1959; Liu et al. 2014; Conner et al. 2015; Underwood et al. 2022). The prevailing question remains as to whether the genes controlling these processes in *Malus* are physically linked together in chromosomal space, although the FCSS results provide indirect insight on this topic. In the facultative apomicts confirmed in the current work, haploid parthenogenesis and B_III_ hybrids (presumably from the pollination of an apomeiotic egg cell) never existed in isolation, implying apomictic components may be commonly inherited together. Still, additional genotypes may need to be screened since it is also unclear whether distantly-related *Malus* species share the same genetic basis for this trait. The sequence data generated here represents some of the most diverse WGS resources for apomictic individuals to date and can be used in future comparative -omics studies to map apomixis genes and describe their evolutionary trajectory in apple. These genes have enormous potential for plant breeding, and characterizing them across diverse angiosperm lineages could mitigate existing obstacles to engineering apomixis in crop species (Vernet et al. 2022; Simon et al. 2025).

## AUTHOR CONTRIBUTIONS AND FUNDING SOURCES

CZG conceptualized, designed, and analyzed results of the sequencing screen. VP conducted and analyzed all FCSS data. BG assisted with material collection and processing. TU and AH supervised experiments and secured funding. All authors participated in writing of the manuscript. CZG is supported by a National Science Foundation Postdoctoral Research Fellowship in Biology, award number 2305693. BG is supported by the USDA Agricultural Research Service Plant Genetic Resources Unit. AH is supported by a National Science Foundation Plant Genome Research Program CAREER (award number 2239530).

## ACKNOWLEDGEMENTS

We would like to especially acknowledge and thank the farm staff managing the USDA germplasm collection in Geneva, NY, including Peter Herzeelle. We are grateful to information technology specialist Rich Johnson at the HudsonAlpha Institute for Biotechnology for cluster maintenance and assistance. Undergraduate researchers instrumental for seed collection and part of the DNA extractions included Lauren Womack, Trinity Tennant, Aislie Casey, and Samantha Johnson. Patricia Sanmartin, Laura Griffin, and Haley Hale provided guidance for library preparation. Paul Doran of Twist Bioscience was essential for library preparation optimization and sequencing strategies. Salina Hall, Fanni Lakatos, and other support staff at Discovery Life Sciences were very helpful for designing and multiplexing sequencing lanes. Several staff members at the Genome Sequencing Center (Ada Stewart, Mike Frizzell) also kindly provided technical expertise for lab and bioinformatic analyses.

## SUPPLEMENTARY TABLE LEGENDS

**Supplementary Table 1**: List of accession classifications as stated in the USDA germplasm database (August 2025), their PI numbers, ploidy, the brand of kit used, the number of embryos sequenced, and the number of seeds analyzed for flow cytometry. Fruiting years the seed was collected are in parentheses. The embryos derived from genotypes highlighted in blue were sequenced at a target depth of 6X for their 2023 seed (up to 96 samples per sequencing lane). All other embryos were sequenced at a target depth of 3X (up to 192 samples per sequencing lane).

**Supplementary Table 2:** Sequencing results between mother and child for 1216 embryos, 90 spike-ins, and 432 simulated embryos (read subsets from maternal genotype’s separately-prepared library). **embryo** = embryo # assigned to a true biological replicate (contains _e #), a simulated embryo from the maternal parent’s dataset (contains _sim), or a spikein of the mother’s DNA also prepared with Twist Bioscience’s 96-plex library kit (contains _a, _b, _c, _d, or _e but no number). **mother** = the genotype a sample is derived from; **het_sites_total** = total number of sites compared between an embryo/sample and its mother when she was heteroallelic (0/1); **het_sites_mismatch** = number of sites mismatching between an embryo/sample and its mother when she was heteroallelic (0/1); **het_match_percentage** = percent of sites matching between an embryo/sample and mother when she was heteroallelic (0/1); **hom_sites_total** = total number of sites compared between an embryo/sample and its mother when she was homoallelic (0/0 or 1/1); **hom_sites_mismatch** = number of sites mismatching between an embryo/sample and its mother when she was homoallelic (0/0 or 1/1); **hom_match_percentage** = percent of sites matching between an embryo/sample and mother when she was homoallelic (0/0 or 1/1**); total_sites** = total number of sites compared between embryo/sample and its mother; **total_mismatch** = total number of sites mismatching between embryo/sample and its mother; **raw_cov** = raw depth of sequencing for that embryo/sample, calculated by: ((# of reads aligned to genome * 150 bp) / 650,000,000 bp); **seed_year** = fruiting year the seed was collected; **quadrant** = quadrant an embryo falls in. Q1 = Selfing or haploid parthenogenesis (rare). Q2 = apomixis or selfed unreduced egg cells (self fertilization of clonal egg). Q3 = sexual reproduction, particularly meiotic reduction, recombination, and foreign pollination; Q4 = foreign pollination with low rates of recombination or foreign pollination of an unreduced egg cell (which would result in a ploidy level increase).

**Supplementary Table 3:** Flow cytometry seed screen (FCSS) results for 26 apple genotypes. Columns “Mean X” record the position of peaks on the x-axis and are supplemented by the coefficient of variance (CV). The ploidy of the embryo and endosperm was subsequently calculated from the obtained peak ratios. Column “Mating system” assigns each seed to one of the following systems: SE = sexual formation, AA = autonomous apomixis, PG = pseudogamy, HP = haploid partenogenesis, GR = genome reduction, GI = genomic increase. The contributions of female (F) and male (P) gametes to seed formation were subsequently calculated from the ploidy levels of the embryo (Emb) and the endosperm (End). For sexual seeds: F = End - Emb, P = Emb - F; for pseudogamous seeds: F = Emb, P = End - 2*Emb and for autonomous apomictic: F = Emb, P = End - 2Emb ; . The last mentioned equation was modified if more than two polar nuclei were involved in endosperm formation in pseudogamous seeds: P = End - N*Emb, where N = number of polar nuclei. U= unreduced gamete; A = apomictic formation; AUP = aneuploidy.

---

**Supplementary Figure 1:**
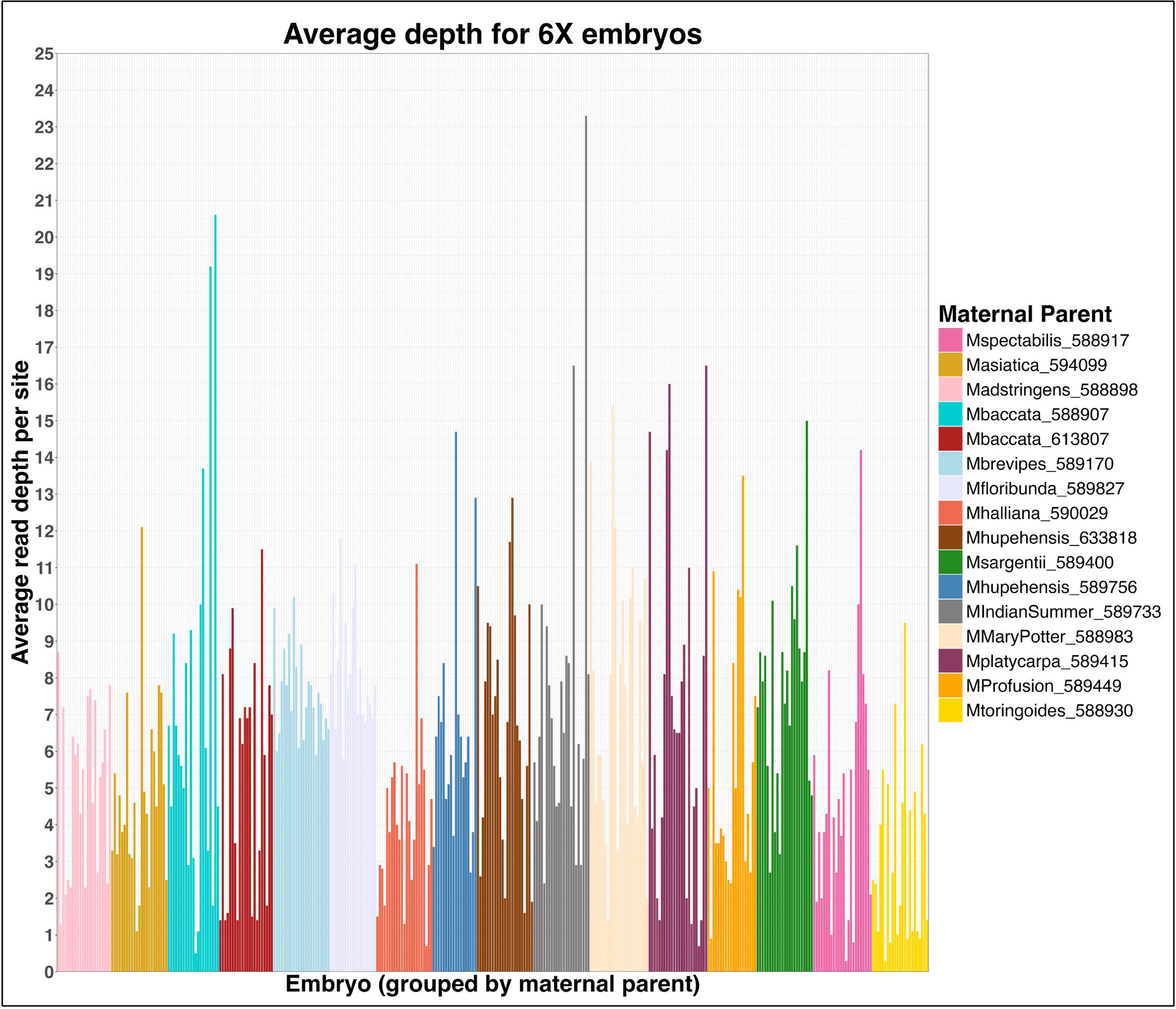
Average depths of embryos sequenced at approximately 6X, grouped by the maternal parent to show the association between depth sequenced and mother, as well as the variation of depth within each mother. For example, *Malus toringoides* 588930 embryos were clearly sequenced at lower depths on average compared to *M. brevipes* 589170 or *M. floribunda* 589827, and these latter two genotypes showed less variable embryo depths compared to other mothers.

**Supplementary Figure 2:**
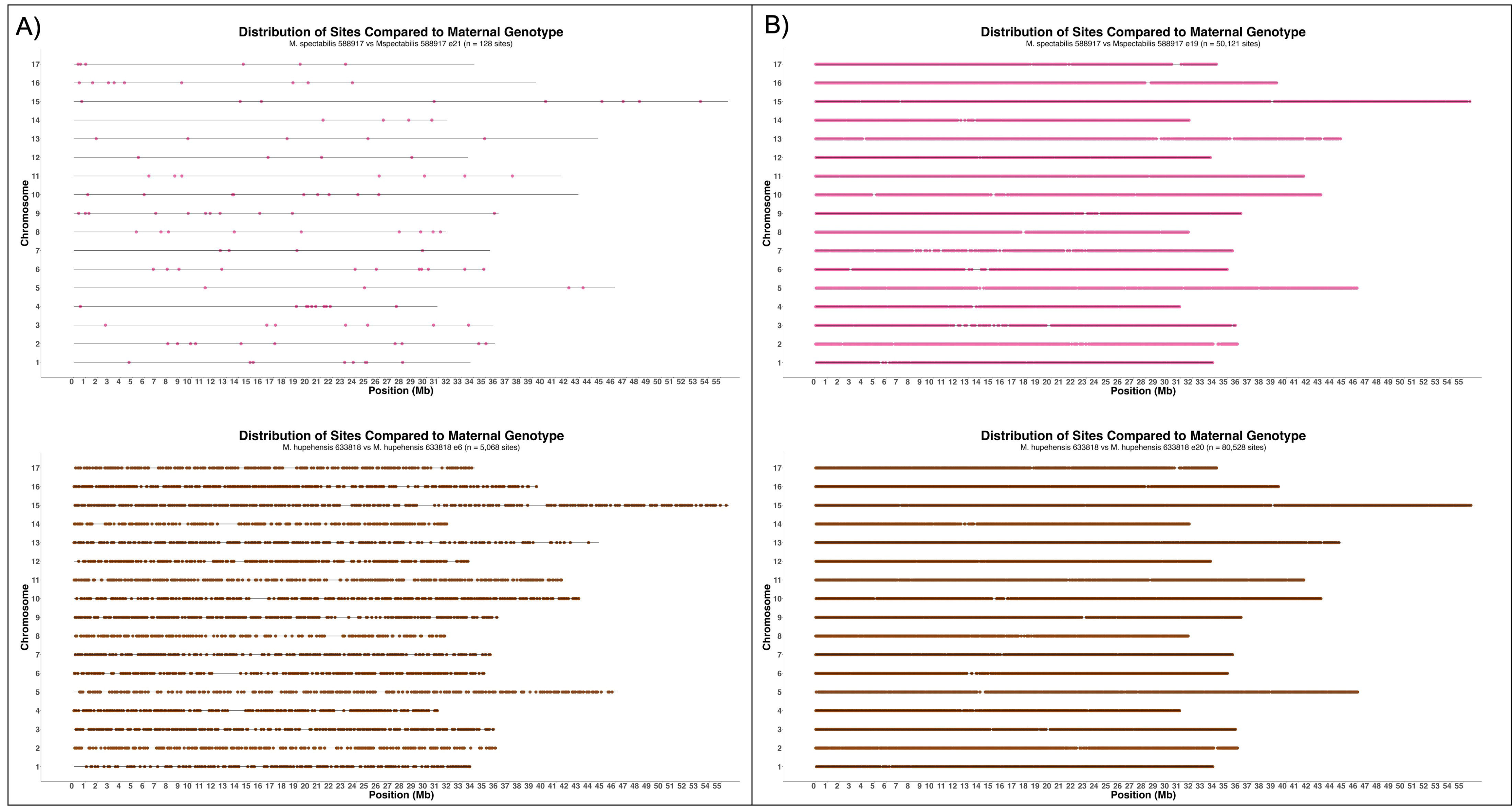
Locations of biallelic sites compared between mother and child for two embryos each from *M. spectabilis* 588917 and *M. hupehensis* 633818. The distributions along the lengths of apple’s 17 chromosomes are shown for **A)** embryos of the lowest depth for each mother and **B)** embryos with an average depth for that mother.

**Supplementary Figure 3:**
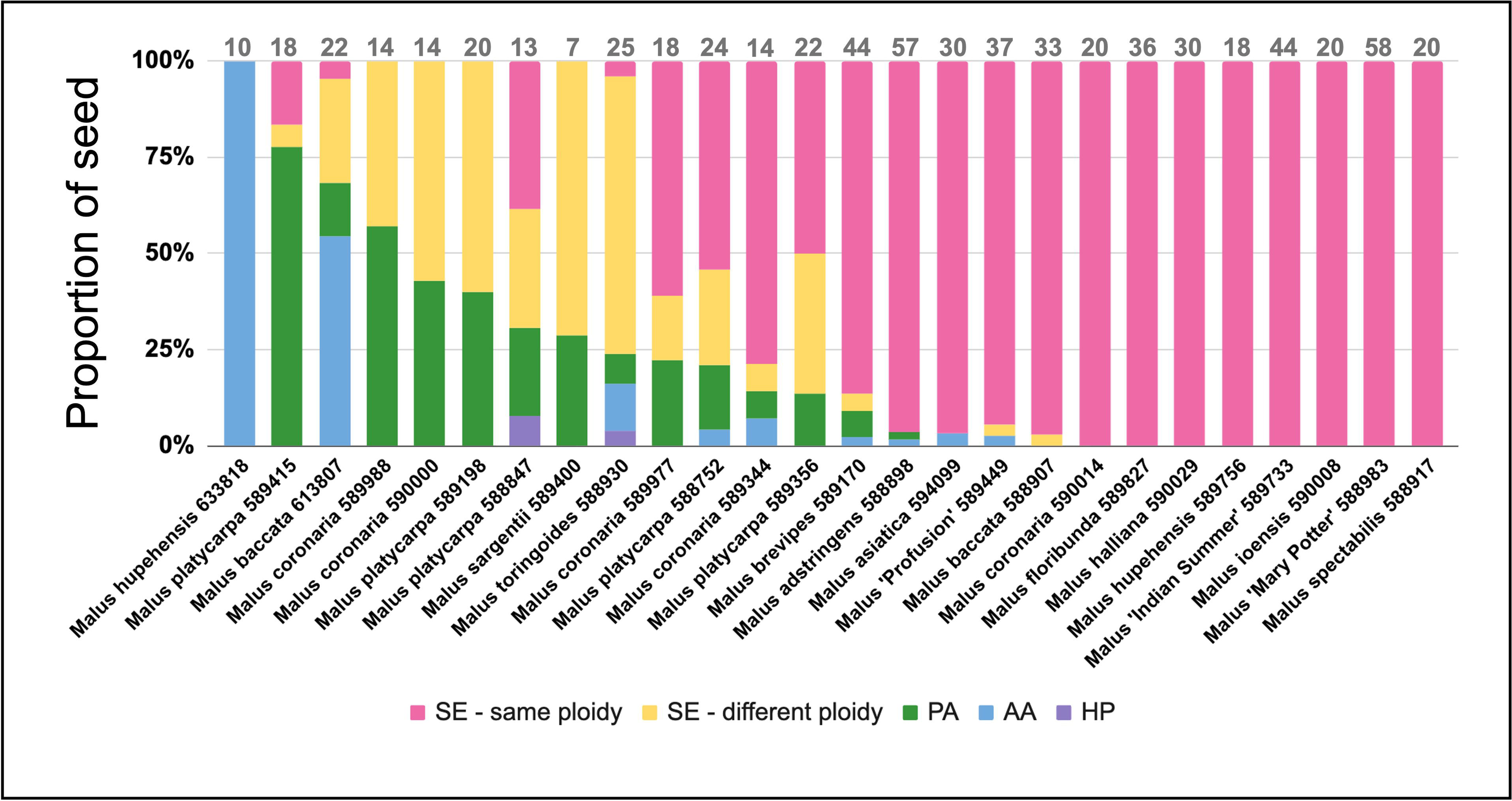
Graphical representation of the FCSS results, displaying the proportion of seed analyzed for each genotype that was determined to be sexually derived (SE) and the same ploidy as the maternal genotype, sexually derived and a different ploidy, a product of pseudogamous apomixis (PA), a product of autonomous apomixis (AA), or a produced by haploid parthenogenesis (HP). Total seeds analyzed per genotype are noted at the top of each respective bar.

**Supplementary Figure 4:**
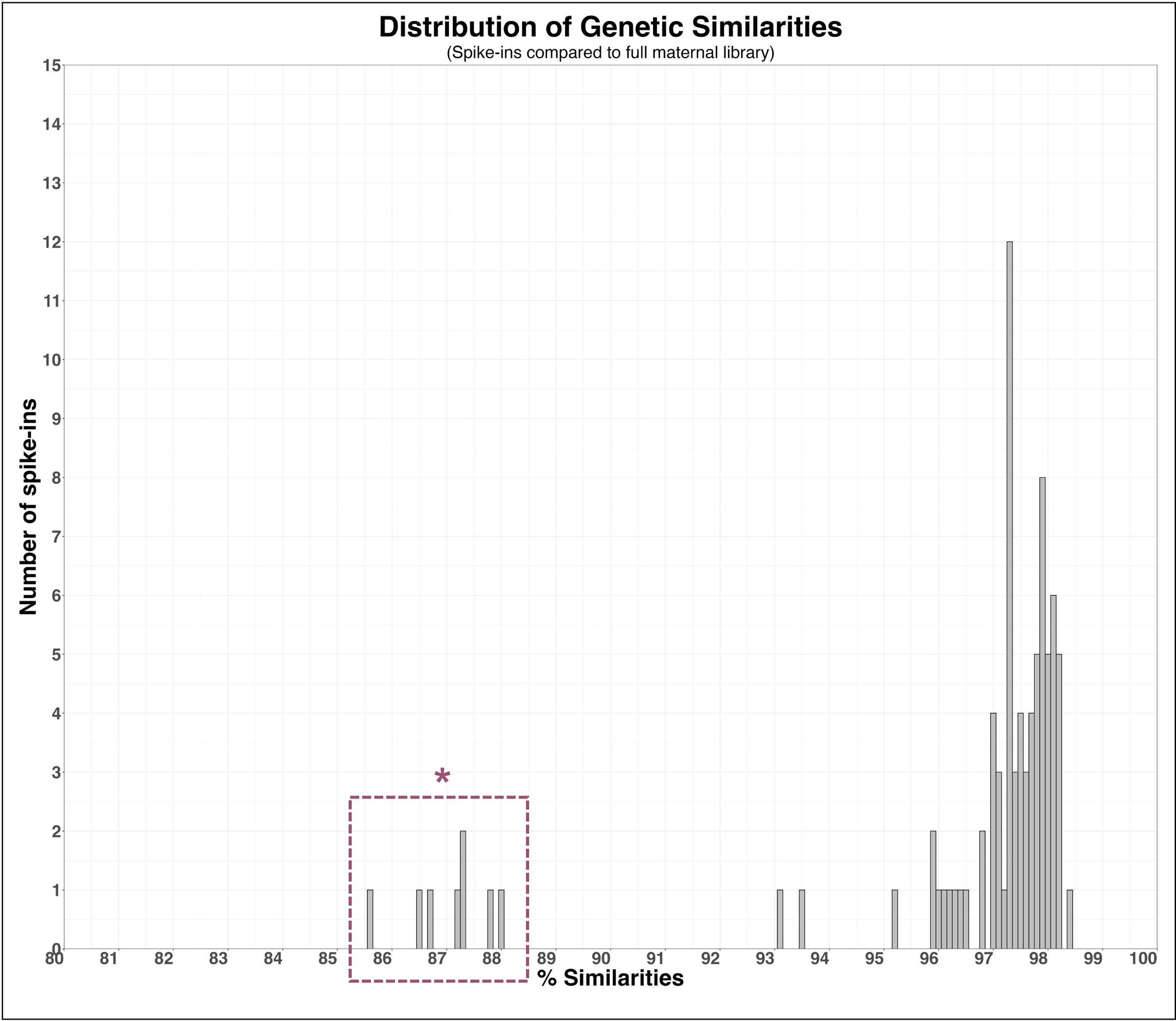
Histogram showing the % similarities between maternal leaf DNA extracts prepared with either a SeqWell or NEBNext library kit and the Twist Bioscience’s 96-plex kit (spike-ins). Spike-ins prepared from tetraploid *Malus platycarpa* genotypes all showed substantially lower similarities to the same DNA prepared with SeqWell or NEBNext. These observations are indicated in purple.

